# Functional angiogenesis requires microenvironmental cues balancing endothelial cell migration and proliferation

**DOI:** 10.1101/868497

**Authors:** William Y. Wang, Daphne Lin, Evan H. Jarman, William J. Polacheck, Brendon M. Baker

## Abstract

Angiogenesis is a complex morphogenetic process that involves intimate interactions between multicellular endothelial structures and their extracellular milieu. *In vitro* models of angiogenesis can aid in reducing the complexity of the *in vivo* microenvironment and provide mechanistic insight into how soluble and physical extracellular matrix cues regulate this process. To investigate how microenvironmental cues regulate angiogenesis and the function of resulting microvasculature, we multiplexed an established angiogenesis-on-a-chip platform that affords higher throughput investigation of 3D endothelial cell sprouting emanating from a parent vessel through defined biochemical gradients and extracellular matrix. We found that two fundamental endothelial cell functions, migration and proliferation, dictate endothelial cell invasion as single cells vs. multicellular sprouts. Microenvironmental cues that elicit excessive migration speed incommensurate with proliferation resulted in microvasculature with poor barrier function and an inability to transport fluid across the microvascular bed. Restoring the balance between migration speed and proliferation rate rescued multicellular sprout invasion, providing a new framework for the design of pro-angiogenic biomaterials that guide functional microvasculature formation for regenerative therapies.

## INTRODUCTION

The microvascular network of arterioles, capillaries, and venules is a critical component of the circulatory system required for the function and maintenance of nearly every tissue in the human body. Once regarded simply as passive fluidic microstructures, it is now understood that the microvasculature dynamically alters its structure and function to service the changing metabolic demands of tissues^1,2^. Rapid changes in microvessel diameter through vasoconstriction or vasodilation allow for temperature and blood pressure regulation^3^. In response to tissue injury, local microvasculature rapidly adjusts permeability, enabling immune cells to extravasate and fight infection; over longer timescales, the microvasculature expands to revascularize the healing tissue^4^. Angiogenesis, the formation of new microvasculature from an existing parent vessel, is the predominant method by which microvasculature extends and is critical to tissue healing and homeostasis^5^. Indeed, dysregulated angiogenesis producing excessive or insufficient microvasculature is a hallmark of many diseases, and as such, microvascular morphology and function are clinical indicators of pathology^6,7^. During cancer progression for example, abnormal gradients of soluble pro-angiogenic factors recruit endothelial cells (ECs) from adjacent tissues to invade into the tumor stroma^8^. This rapid and excessive angiogenesis results in a high density of disorganized and highly permeable neovessels that facilitates tumor growth and provides metastatic access^9^. In contrast, insufficient angiogenesis impairs tissue regeneration for example in cardiac ischemia or diabetic foot ulcers^10^. An understanding of how the surrounding microenvironment appropriately guides the formation of functional microvasculature rather than excessive or insufficient angiogenesis in disease contexts would be critical to designing vascularized biomaterials for regenerative medicine and novel therapies to treat vasculopathies^6,11,12^.

Observations consistent across a wide range of *in vivo* and *in vitro* models of angiogenesis have established several key steps including (1) chemokine gradients promoting endothelial tip cell formation and directed invasion into the extracellular matrix (ECM), (2) collective migration of leading tip cells and ensuing stalk cells, (3) proliferation and lumenization of the invading strand, and (4) maturation into functional neovasculature^13,14^. Each of these steps is regulated by both biochemical and physical microenvironmental cues presented by the surrounding ECM, the 3D fibrous, collagenous meshwork through which EC sprouts navigate^15^. As characterizing and tuning microenvironmental cues (e.g. profiling biochemical gradients and controlling physical ECM properties) is challenging *in vivo, in vitro* models have proven instrumental in providing mechanistic insight into each step of angiogenesis^16^. To build our understanding of how ECs migrate collectively, 2D scratch wound assays are widely utilized. While these assays have provided detailed insight into the molecular pathways governing collective migration of EC monolayers^17^, the model fails to recapitulate the 3D nature of sprouting morphogenesis^18^. EC outgrowth assays from spheroids or microbead carriers embedded within 3D ECM have been instrumental in studying the role of matrix proteolysis and tip vs. stalk cell identity and dynamics^19–21^. However, sprouts in these models do not originate from an accessible lumenized parent vessel, making it difficult to assess key microvascular functions such as fluidic connectivity and permeability. More recently, advances in biomicrofluidics and efforts to engineer tissues-on-chips have generated 3D human engineered microvessels^22–24^; these models have been utilized to study how chemokine gradients, ECM degradability, shear stress and support cells regulate vessel barrier function, EC sprouting and tumor and immune cell extravasation^22,23,25–27^. While much information has been learned from these various models, how EC migration speed and proliferation – two fundamental cell processes required for angiogenesis – influence the formation and function of subsequent microvasculature remains unresolved.

In this work, we multiplexed a microfluidic device that recapitulates key aspects of sprouting morphogenesis, namely the directional, chemokine-driven invasion of ECs from the stable and quiescent endothelium of a fluid-bearing arteriole-scale parent vessel into the surrounding 3D ECM. We used this biomimetic model to investigate how soluble and physical ECM properties regulates EC invasion speed and proliferation during sprouting and how independently tuning the rate of these two basic cell functions influences the quality of formed microvasculature. We find that the formation of functional microvasculature capable of transporting fluid and performing barrier function requires a delicate balance between EC migration speed and proliferation rate. Furthermore, we demonstrate that aberrant angiogenic sprouting driven by altered physical and soluble microenvironmental cues can in fact be rescued by correcting the imbalance between these two fundamental EC functions. As the proper vascularization of large tissue engineered constructs remains an outstanding challenge for the biomedical engineering community, the findings of this work establish a new framework for biomaterial design parameters that balance EC migration speed and proliferation to optimally generate functional microvasculature.

## RESULTS

### Multiplexed angiogenesis-on-a-chip platform

To investigate how soluble and physical microenvironmental cues regulate angiogenic sprouting, we adapted a previously established microfluidic device that recapitulates 3D EC sprouting morphogenesis from an arteriole-scale parent vessel^26,27^. We improved the fabrication throughput of these devices by reducing their assembly to a single layer design in addition to multiplexing the number of devices resulting from each fabrication such that a single chip contains a 2×4 array of devices (Fig. 1a). To generate parent vessels in these devices, a hydrogel precursor solution is first injected through device ports and is allowed to crosslink around two parallel needles (300 µm diameter) suspended across each device’s central chamber (Fig. 1a-b). The void space created after needle extraction forms a pair of parallel hollow channels fully embedded within extracellular matrix (ECM) terminating in two media reservoirs (Fig. 1b). ECs seeded into one channel of each device attach to the inner channel surface and self-assemble into a perfused endothelialized tube serving as the parent vessel (Fig. 1c-e). Within 24 hours, the assembled EC monolayer of the engineered parent vessel localizes VE-cadherin to cell-cell junctions, and maintains a consistent diameter and cell density over a range of collagen densities (Fig. 1d and Supplemental Fig. 1a-d).

**Figure 1.**
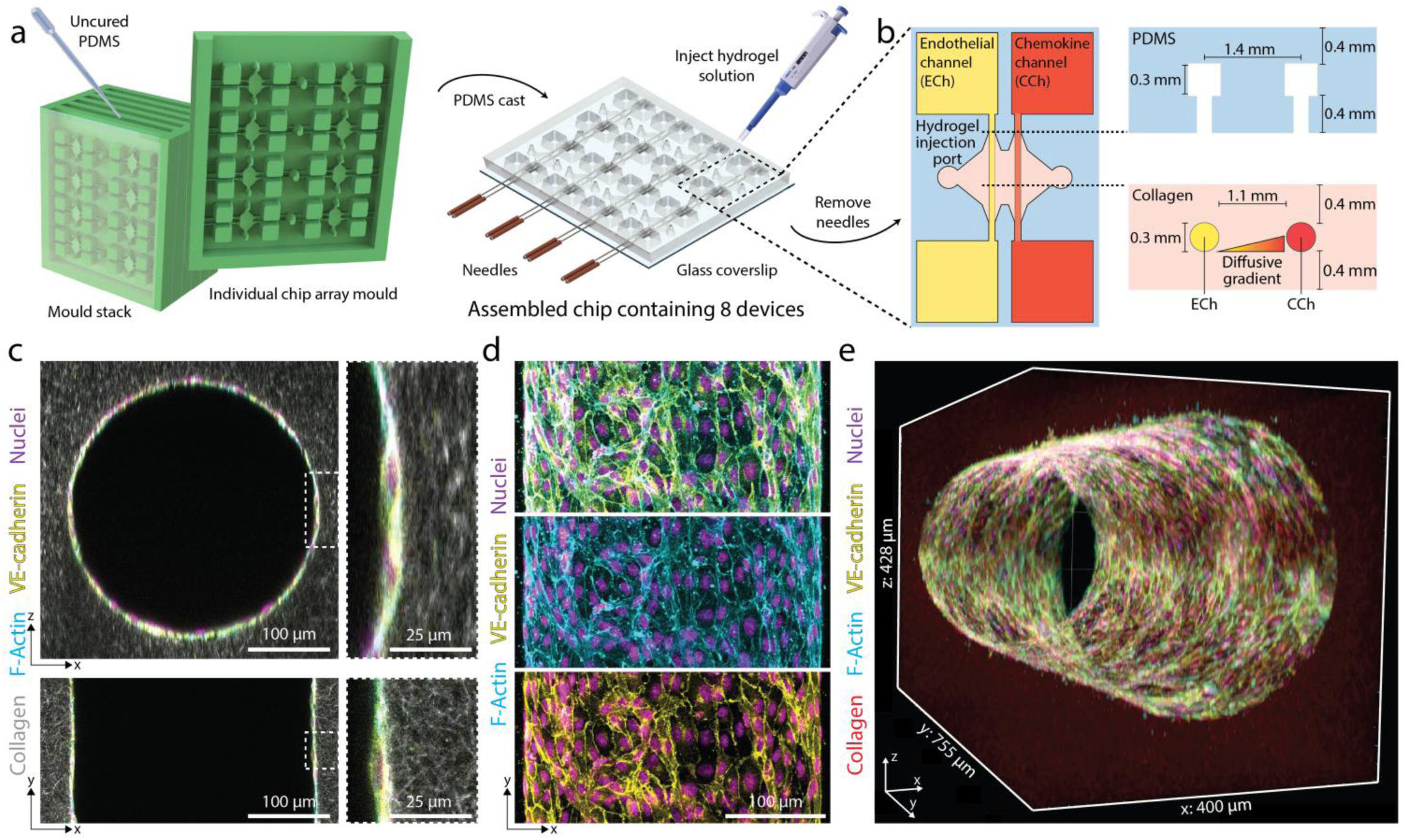
Multiplexed angiogenesis-on-a-chip platform. **a**, PDMS replica casts from 3D-printed moulds are bonded to glass coverslips. Each chip is composed of a 2×4 array of single devices. A pair of needles are inserted into each device, and type I collagen solution is injected into each device and gelled around needles. Hollow channels are generated upon needle removal. **b**, Inserted needles are suspended above the glass coverslip bottom and below the PDMS housing to form 3D channels fully embedded within a collagen hydrogel. Each device is composed of two parallel channels. The endothelial channel (ECh) is seeded with endothelial cells to form the parent vessel. Pro-angiogenic factors are added to the chemokine channel (CCh) and form a diffusive gradient to promote 3D endothelial cell invasion across the collagen hydrogel. **c**, Representative images of x-z (top) and x-y (bottom) orthogonal views of parent vessels formed within fluorescently labeled 3 mg ml^−1^ collagen hydrogel 24-hours post EC seeding. Insets indicated with dashed white lines. **d**, Representative images of x-y (max intensity projection) formed within a 3 mg ml^−1^ collagen hydrogel 24-hours post EC seeding. **e**, 3D rendering of parent vessel formed within fluorescently labeled 3mg ml^−1^ collagen hydrogel.

### Soluble factors regulate multicellular sprouting

Utilizing 3 mg ml^−1^ collagen, parent vessels cultured in EGM2 proved stable as single ECs minimally invaded (5.5 ± 14.9 µm) into the ECM over 5-day culture under continual reciprocating flow (Fig. 2a, c). To induce EC invasion, we introduced an established EC chemoattractant, sphingosine 1-phosphate (S1P), to the adjacent chemokine channel to produce a diffusive gradient that drives directional 3D EC invasion through the ECM^26–28^. We found EC invasion depth over 5-day culture to be dependent on [S1P], with increasing [S1P] resulting in increased invasion speed (Fig. 2a, c). To assess the morphologic quality of EC invasion, we categorized ECs as isolated single cells or multicellular sprouts and determined the ratio of sprouts to single cells as a metric of invasion multicellularity. Due to variations in invasion depth across conditions, we restricted quantification of single cells to 150 µm from the leading invasive front and defined sprouts as contiguous multicellular structures with a length greater than half the max invasion depth (Supplemental Fig. 2). Although [S1P] clearly mediated cell invasion in a dose-dependent manner, the phenotype of invading ECs was primarily as single, disconnected cells and multicellular sprouts were rarely observed (Fig. 2a, c, e-g). Due to the low levels of EC proliferation observed in these conditions (Fig. 2d) and given previous evidence that EC proliferation is required for angiogenesis *in vivo*^29,30^, we hypothesized that enhancing proliferation rates would increase the number of ECs collectively invading as multicellular sprouts. Media supplementation with 25 ng ml^−1^ phorbol 12-myristate 13-acetate (PMA), another well-established pro-angiogenic factor and potent activator of PKC^26,27,31,32^, resulted in elevated proliferation rates as assayed by EdU incorporation (Fig. 2d, Supplemental Fig. 2). Few invading ECs were observed at 0 nM S1P with 25 ng ml^−1^ PMA implying that the addition of PMA alone does not induce EC invasion (Fig. 2c). Invasion speed remained S1P dose-dependent in the presence of 25 ng ml^−1^ PMA while proliferation rates proved independent of S1P dose (Fig. 2c-d). In support of our hypothesis, PMA supplementation and elevated EC proliferation corresponded to significant increases in the number of invading multicellular sprouts at each level of [S1P] (Fig. 2d-g). In conditions with 25 ng ml^−1^ PMA, sprout invasion depth over 5-day culture anti-correlated with the ratio of multicellular sprouts to single ECs (Fig. 2c, g); the highest level of S1P (500 nM) resulting in the fastest invasion speed, most single cells, and fewest multicellular sprouts (Fig. 2c, e-f). These data clearly indicate proliferation in this model is a requirement for multicellular sprout invasion, supporting previous observations *in vivo*^29,30^. Furthermore, while a chemokine gradient is an additional key requirement for angiogenesis, stronger gradients that increase invasion speed elicit a single cell migration phenotype in lieu of multicellular sprouts.

**Figure 2.**
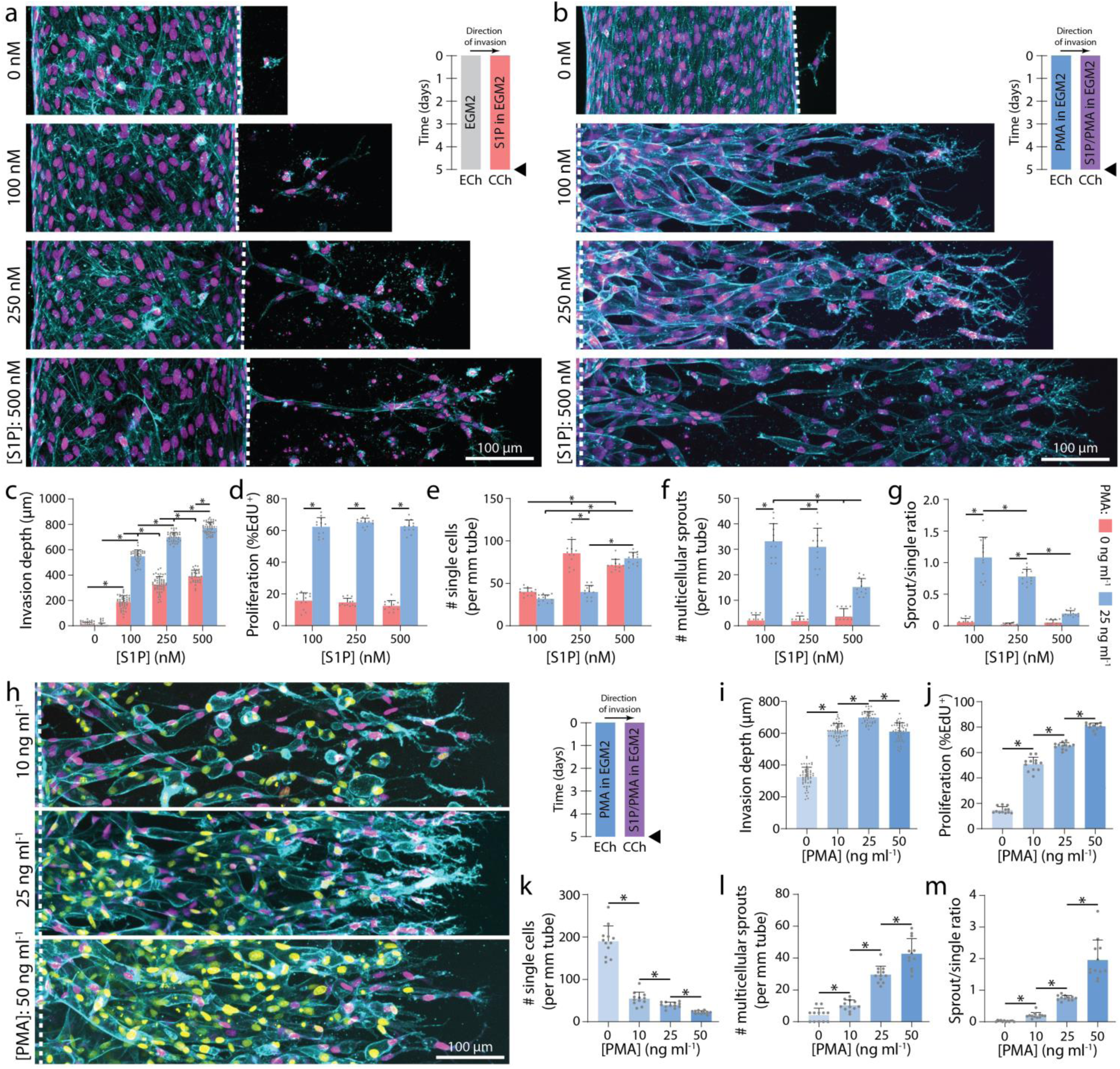
Soluble factors modulate EC invasion, proliferation, and multicellular sprouting. **a**, Representative images (max intensity projection) of invading endothelial cells in response to varying [S1P]. **b**, Representative images (max intensity projection) of invading endothelial cells in response to varying [S1P] with 25 ng ml^−1^ PMA. F-actin (cyan), nucleus (magenta), and dashed white lines indicate parent vessel edge (**a-b**). **c-g**, Quantifications of invasion depth, proliferation, and morphology of invading endothelial cells as single cells or multicellular sprouts. **h**, Representative images (max intensity projection) of invading endothelial cells in response to varying [PMA] with 250 nM S1P. **i-m**, Quantifications of invasion depth, proliferation, and morphology of invading endothelial cells as single cells or multicellular sprouts. UEA (cyan), nucleus (magenta), EdU (yellow). All data presented as mean ± s.d.; * indicates a statically significant comparison with P<0.05 (one-way analysis of variance). For invasion depth analysis (**c, i**), n≥32 vessel segments (each 100 µm length) per condition; for proliferation and migration mode analysis (**d-g, j-m**), n=12 vessel segments (each 800 µm length) per condition.

To investigate the effects of proliferation rates on multicellular angiogenic sprouting, we next varied [PMA] while maintaining [S1P] constant at 250 nM. Proliferation rates proved PMA dose-dependent and positively correlated with the number of multicellular sprouts (Fig. 2h, j-m). Although PMA alone did not induce EC invasion (Fig. 2b-c), increasing [PMA] in the presence of S1P resulted in a biphasic relationship with invasion depth (Fig. 2i). The influence of increasing [PMA] from 0 to 25 ng ml^−1^ on increased invasion depth may be due to a greater number of invading cells each secreting matrix metalloproteinases (MMPs). Elevated MMP levels would hasten localized matrix degradation, allowing ECs to more rapidly generate open space within 3D ECM required to advance forward. Interestingly, at the highest tested concentration of PMA resulting in the most proliferation, invasion depth decreased (Fig. 2i). As cells are transiently non-migratory during mitosis, this decrease may stem from frequent proliferative events hampering efficient migration of ECs. Varying [PMA] and thus PKC activation may have multiple downstream effects, so we treated ECs with mitomycin C (a crosslinker that prevents DNA replication and inhibits mitosis) to confirm the role of proliferation as the primary effector of enhancing multicellular sprout invasion. Even in the presence of the highest level of PMA, proliferation-inhibited ECs invaded only as single cells with significantly decreased invasion depth as compared to controls (Supplemental Fig. 3a-f). These studies therefore indicate cell proliferation is not only required for multicellular sprout invasion, but additionally influences the rate at which sprouts traverse 3D ECM.

In the experiments above individually modulating migration speed and proliferation, higher migration speeds resulted in disconnected single EC invasion while increasing proliferation rates enhanced multicellular sprouting. We thus hypothesized that proliferation rate commensurate with invasion speed is essential to the collective invasion of multicellular sprouts. Indeed, proportionally increasing or decreasing both [S1P] and [PMA] simultaneously resulted in invasion depth and proliferation rate increases or decreases, respectively, but did not alter the ratio of multicellular sprouts to single ECs (Fig. 3a-g). This suggests that multicellular angiogenic sprouting requires a critical balance between EC invasion speeds and proliferation rates, which can be finely tuned by these two established soluble pro-angiogenic factors. While balanced soluble conditions maintained similar invasion multicellularity, the magnitude of these cues influenced sprout diameter, with higher levels of S1P and PMA increasing sprout diameter (Fig. 3a-b, h). Larger neovessel diameters would allow for increased fluid transport, but smaller diameter vessels would allow for increased capillary density and more efficient nutrient-waste exchange. Although this requires further investigation, here we demonstrate control of neovessel diameter over the 5-25 µm range of capillary diameters reported *in vivo*^2,33^.

**Figure 3.**
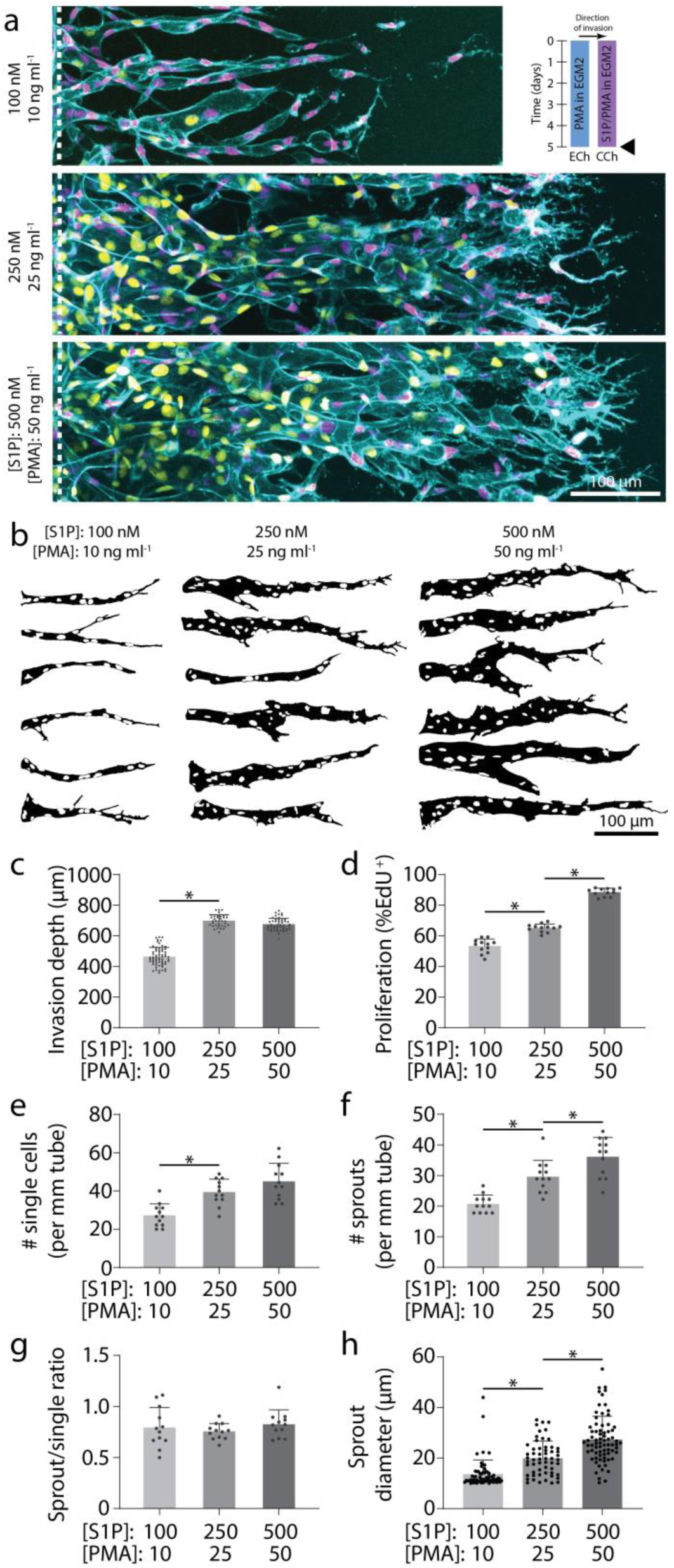
Multicellular sprouting requires balanced migration and proliferation. **a**, Representative images (max intensity projection) of invading endothelial cells in response to proportional variations of [S1P] and [PMA]. UEA (cyan), nucleus (magenta), EdU (yellow). **b**, Representative sprout outlines from conditions in (**a**). **c-h**, Quantifications of invasion depth, proliferation, morphology of invading endothelial cells as single cells or multicellular sprouts, and sprout diameter. All data presented as mean ± s.d.; * indicates a statically significant comparison with P<0.05 (one-way analysis of variance). For invasion depth analysis (**c**), n≥36 vessel segments (each 100 µm length) per condition. For proliferation and migration mode analysis (**d-g**), n=12 vessel segments (each 800 µm length) per condition. For sprout diameter analysis (**h**), n≥56 sprout segments.

### Differential roles of tip and stalk endothelial cells

Previous work indicates that tip cells responsive to angiogenic chemokine gradients lead invading sprouts by degrading the ECM and guiding ensuing stalk cells^13,14^. To investigate whether tip cells require chemokine receptors, we performed mosaic sprouting studies where untreated ECs were mixed with GFP-labeled ECs pre-treated with FTY720 (FTY720-GFP-ECs), an S1P receptor inhibitor. Preliminary studies confirmed the inhibitory effect of FTY720, as ECs pre-treated with 100 nM FTY720 prior to device seeding demonstrated minimal invasion over 5-day culture despite appropriate soluble conditions of 250 nM S1P and 25 ng ml^−1^ PMA (S250:P25) (Supplemental Fig. 4a-b). Furthermore, when ECs were allowed to sprout for 3 days prior to FTY720 treatment, subsequent S1P receptor inhibition halted any further advance of already established sprouts (Supplemental Fig. 4a-b). Performing mosaic studies with controlled ratios of FTY720-GFP-ECs vs. untreated ECs, a higher fraction of FTY720-GFP-ECs decreased invasion depth, the number of single ECs, and thereby enhanced sprout multicellularity as evident by increased sprout-single cell ratios (Fig. 4a, e-i). Furthermore, FTY720-GFP-ECs primarily remained in the parent vessel and did not assume the tip cell position of invading sprouts. FTY720-GFP-ECs were occasionally found within sprout stalks, suggesting that pushing or pulling forces from adjacent stalk cells may enable the movement of FTY-GFP-ECs lacking functional S1P receptors.

**Figure 4.**
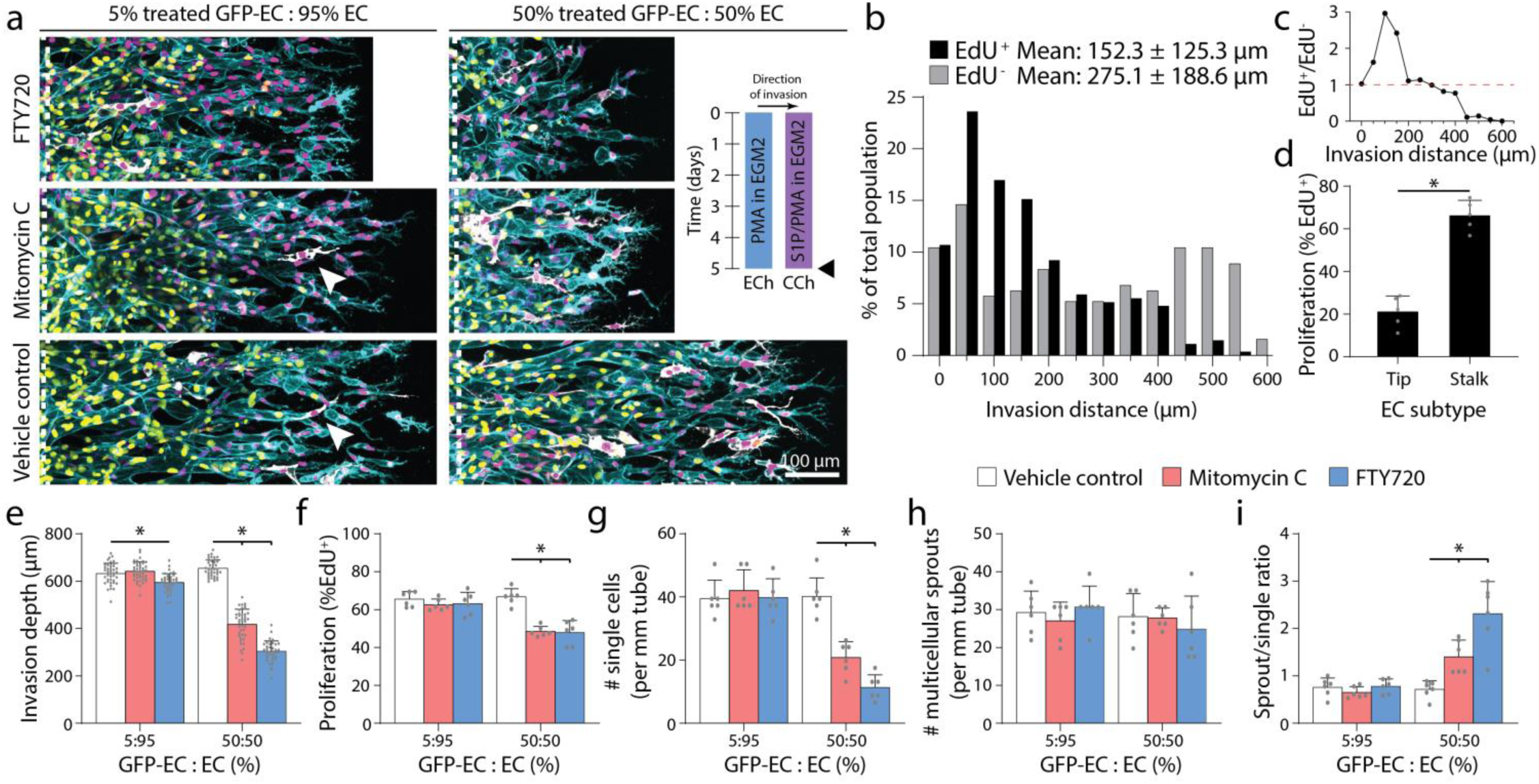
Invading tip cells require chemokine receptors but do not require proliferative capacity. **a**, Representative images (max intensity projection) of invading endothelial cells with S250:P25 and varying ratios of treated GFP-EC to untreated EC with indicated treatment. UEA (cyan), nucleus (magenta), EdU (yellow), GFP-EC (white). **b**, Histogram of EdU^+^ and EdU^−^ invaded endothelial cells with S250-P25 after 5-day culture. **c**, Ratio of EdU^+^ to EdU^−^ endothelial cells from (**b**). **d**, Percentage of EdU^+^ tip or stalk cells from (**b**). **e-i**, Quantifications of invasion depth, proliferation, and morphology of invading endothelial cells as single cells or multicellular sprouts. All data presented as mean ± s.d.; *indicates a statically significant comparison with P<0.05 (two-tailed Student’s t-test (**d**) and one-way analysis of variance (**e-i**)). For spatial EdU analysis (**b-c**), n≥192 cells. For tip vs. stalk proliferation analysis (**d**), n= 5 vessel segments (each 800 µm length). For invasion depth analysis (**e**), n≥37 vessel segments (each 100 µm length) per condition. For proliferation and migration mode analysis (**f-i**), n=6 vessel segments (each 800 µm length) per condition.

Previous work *in vivo* has shown that the frequency of proliferation is spatially segregated by EC subtype (i.e. tip and stalk ECs)^19,30^. Examining the localization of proliferation during sprouting angiogenesis in this model, ECs at the invasion front were indeed the least proliferative (Fig. 4b-c). Furthermore, while only a small percentage of tip ECs underwent proliferation, the majority of EdU^+^ nuclei were positioned within sprout stalks closest to the parent channel (Fig. 4b-d). To test whether tip cells require proliferative capacity, we performed additional mosaic studies seeding parent vessels with mixtures of mitomycin C pre-treated GFP-labeled ECs (MitoC-GFP-EC) and untreated ECs. At low mosaic ratios (5% MitoC-GFP-EC) where overall sprouting was not influenced by non-proliferating ECs, MitoC-GFP-ECs were observed at the tip cell position, confirming previous observations *in vivo* that tip cells do not require the capacity to proliferate (Fig. 4a, e-i)^19,30^. At high mosaic ratios (50% MitoC-GFP-EC), overall proliferation rates decreased as expected (Fig. 4f). While impaired proliferation should decrease multicellular sprouting based on the studies above (Fig. 2h-m), invasion speeds also decreased such that the balance between migration speed and proliferation was maintained and the number of multicellular sprouts did not differ from controls (Fig. 4e-f, h). With decreased invasion speed, fewer tip cells at the invasive front broke away as single cells resulting in enhanced ratios of sprouts to single cells (Fig. 4g-i). Taken together, tip and stalk cells perform differential roles during sprouting and possess distinct requirements for chemotaxis and proliferation. Tip cells require chemokine receptors to migrate in response to soluble gradients, but do not require proliferative capacity; in contrast, proliferation primarily occurs in ensuing stalk cells, perhaps providing a requisite cell density needed to maintain intercellular connectivity within the invading multicellular structure.

### Functional assessment of fluidic connectivity and permeability

A critical function of microvasculature is the transport of nutrients, waste, platelets, and immune cells. To assess the patency and fluidic connectivity of neovessels, we allowed ECs to fully traverse the 1.1 mm wide collagen matrix and reach the adjacent CCh. Time-lapse confocal imaging while introducing fluorescent microspheres (Ø = 1 µm) into ECh reservoirs enabled rapid assessment of flow across the formed vascular bed (ECs spanning ECM). In vascular beds generated under balanced levels of invasion speed and proliferation rates (i.e. conditions resulting in high sprout-single ratios), beads readily flowed through the neovessel network spanning across ECh to CCh demonstrating fluidic connectivity of functional microvasculature (Fig. 5a). In stark contrast, dysfunctional microvascular beds formed with excessively high invasion speed (S500:P25) or deficient proliferation (S250:P10) resulting in disconnected sprouts (i.e. low sprout-single ratio), fluorescent microspheres failed to flow across the ECM space and remained sequestered near the endothelialized parent channel (Fig. 5a). Indeed, max intensity projections of the full height of the 3D vascularized ECM revealed a complete absence of bead flow, confirming observations from individual z-slices captured during time-lapse imaging (Fig. 5b).

**Figure 5.**
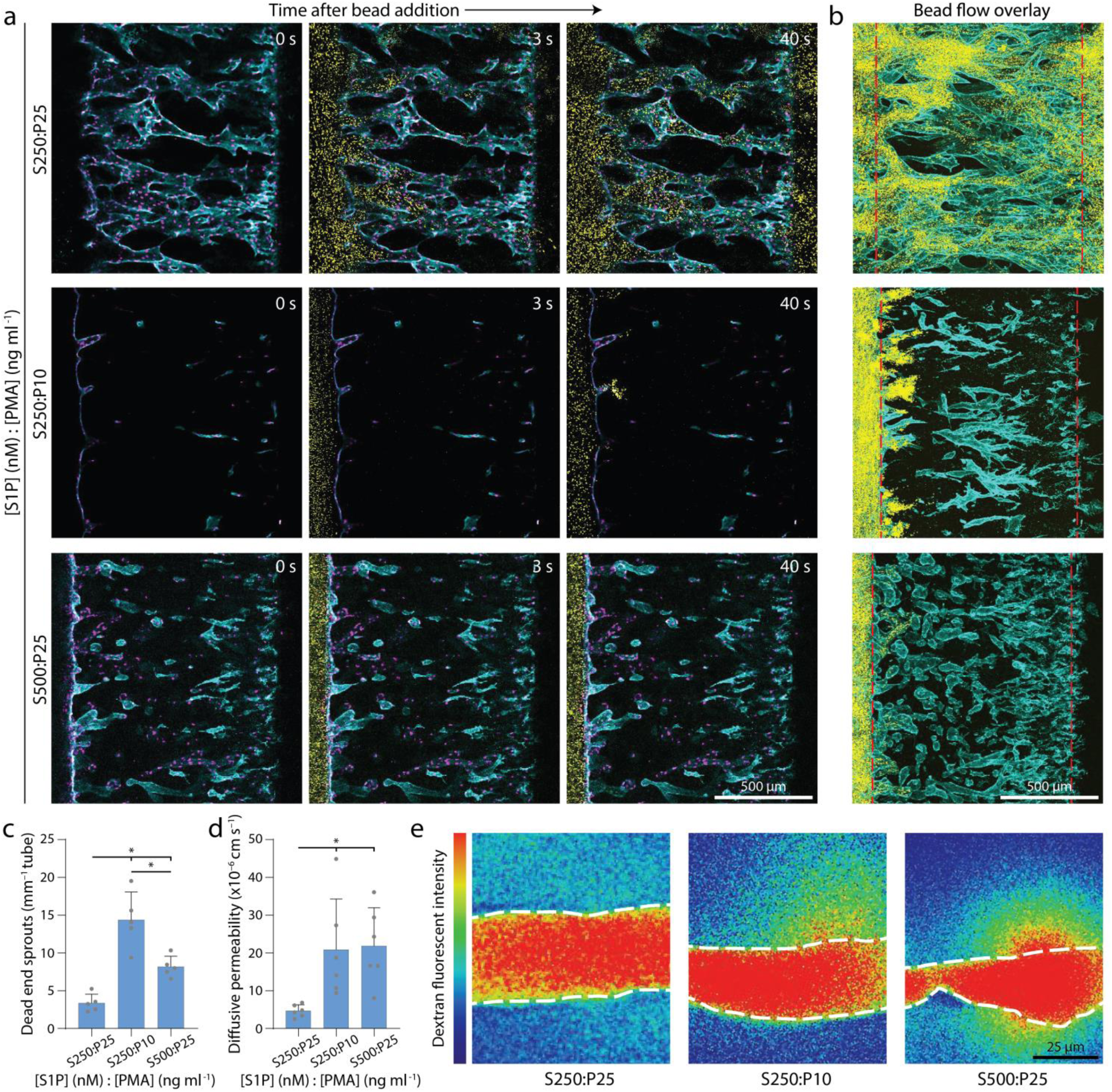
Balanced migration and proliferation optimize microvasculature fluidic connectivity and permeability. **a**, Representative time course images of 1 µm bead flow across vascular beds formed with balanced cues (top), low proliferation (middle), and high migration (bottom). Nucleus (magenta), F-actin (cyan), beads (yellow). **b**, Max intensity projection at steady state from (**a**); red dashed lines indicate ECh and CCh edges. F-actin (cyan), beads (yellow). **c**, Quantification of sprouts incapable of fluidically transporting beads. **d**, Quantification of diffusive permeability of 70-kDa dextran. **e**, Representative heat map images of 70 kDa dextran diffusion across neovessels within vascular beds formed with balanced cues (left), low proliferation (center), and high migration (right). White dashed lines indicate vessel edge. All data presented as mean ± s.d.; * indicates a statically significant comparison with P<0.05 (one-way analysis of variance). n=5 devices per condition (**c**), n=6 neovessels per condition (**d**).

Barrier function and endothelial permeability regulated by cell-cell junctions are a closely related aspect of microvascular function^23^. To assess the permeability of formed vascular beds, we introduced fluorescent 70 kDa dextran to the endothelial parent channel after ECs fully traversed the collagen matrix reaching the adjacent CCh, allowed flow across the vascular bed, and time-lapse imaged dextran diffusion across neovessel walls into the surrounding ECM. Low permeability was only achieved when vascular beds were formed under soluble cues that balanced invasion speed with proliferation rates (Fig. 5d-e), and are comparable to reported values *in vivo* of protein diffusion across capillaries (4.3×10^−6^ cm s^−1^)^34^. Interestingly, in both bead flow and permeability assessments, imbalanced soluble conditions resulting in disconnected sprouts were composed of neovessels that were highly permeable at the tip cell position, such that 1.0 µm beads collected in the adjacent ECM (Fig. 5c). Taken together, multicellular invasion driven by soluble cues that balance invasion and proliferation yields fluidically functional microvascular beds with low permeability.

### Matrix density regulates sprouting speed and morphology

In addition to soluble cues, physical properties of the ECM are known to regulate EC morphology and function during angiogenesis^15^. We hypothesized that independent of soluble cues, physical ECM cues may also influence the balance between EC migration speed and proliferation, and therefore invasion as multicellular sprouts. For example, ECM density, stiffness, and degradability define the susceptibility of ECM to proteolytic activity required for 3D migration^21,27^. To investigate whether physical properties of ECM influence EC invasion as multicellular sprouts we tuned ECM density by modulating collagen concentration (all previous studies were performed in 3 mg ml^−1^ collagen). Maintaining constant soluble cues of S250:P25, increasing collagen density resulted in decreased invasion depth over 3-day culture (Fig. 6a, d). Interestingly, varying collagen density from 2-6 mg ml^−1^ did not significantly alter EC proliferation rates, perhaps due to PMA’s potent enhancement of proliferation (Fig. 6e). Mimicking the response of EC invasion speed to [S1P] (Fig. 2c), decreasing matrix density increased EC invasion speeds with a parallel shift in the morphology of invading ECs from primarily multicellular sprouts towards single cells (Fig. 6d, f-h). Time-lapse imaging capturing the dynamics of EC invasion revealed that tip ECs break away from the parent vessel more frequently in matrices with low collagen density (Fig. 6b-c).

**Figure 6.**
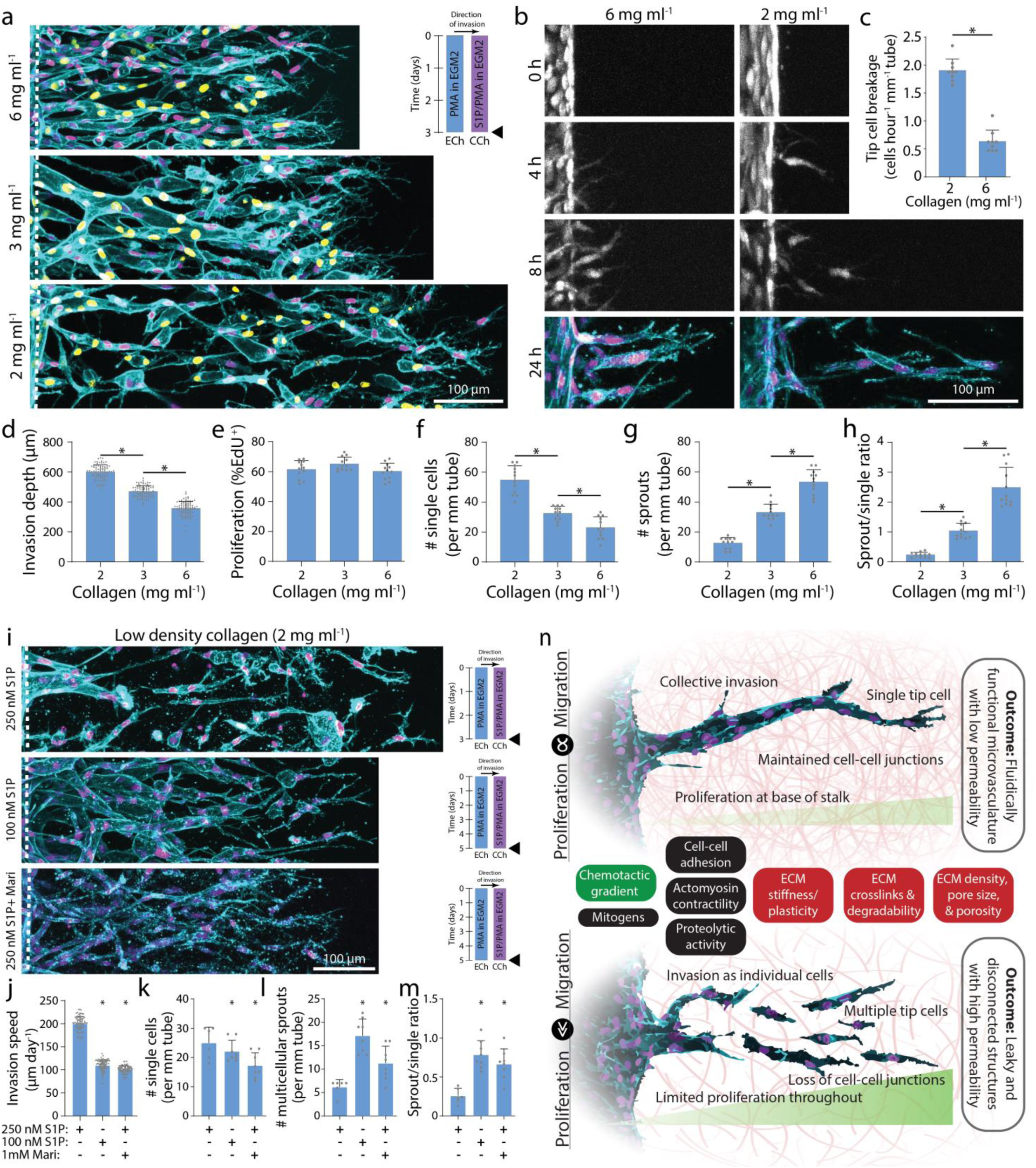
Increasing collagen density decreases invasion speed and enhances multicellular sprouting. **a**, Representative images (max intensity projection) of invading endothelial cells in response to varying collagen density with S250:P25. UEA (cyan), nucleus (magenta), EdU (yellow). **b**, Representative time course images (max intensity projection) of invading endothelial cells (labeled with cell tracker dye) in response to varying collagen density with S250:P25. F-actin (cyan) and nucleus (magenta). **c**, Quantification of the frequency of tip cell breakage events. **d-h**, Quantifications of invasion depth, proliferation, and morphology of invading endothelial cells as single cells or multicellular sprouts from conditions in (**a**). **i**, Representative images (max intensity projection) of invading endothelial cells in 2 mg ml^−1^ collagen with control (S250:P25), decreased S1P (S100:P25), and MMP inhibition (S250:P25). F-actin (cyan) and nucleus (magenta). **j-m**, Quantifications of invasion speed, and morphology of invading endothelial cells as single cells or multicellular sprouts. All data presented as mean ± s.d.; * indicates a statically significant comparison with P<0.05 (two-tailed Student’s t-test (**c, j-m**) and one-way analysis of variance (**d-h**)). For tip cell breakage analysis (**c**), n=9 vessel segments (each 400 µm length) per condition. For invasion depth analysis (**d**), n≥80 vessel segments (each 100 µm length) per condition. For proliferation and migration mode analysis (**e-h, k-m**), n≥6 vessel segments (each 800 µm length) per condition. For invasion speed analysis (**j**), n≥81 vessel segments (each 100 µm length) per condition. **n**, Schematic illustration highlighting the relationship between endothelial cell migration and proliferation on invasion morphology as multicellular sprouts (top; balanced migration and proliferation) vs single cells (bottom; excessive migration or insufficient proliferation). Soluble (green) and physical extracellular matrix (red) cues influence this balance in addition to other potential cell functions (black) that may regulate multicellular sprouting and formation of functional microvasculature.

We thus hypothesized that reducing EC invasion speed in low density collagen would rescue multicellular sprout invasion by allowing trailing stalk cells to maintain intercellular connectivity with leading tip cells. We first decreased EC invasion speed by reducing the strength of the chemokine gradient. Decreasing [S1P] from 250 nM to 100 nM resulted in decreased invasion speed and increased EC invasion as multicellular sprouts (Fig. 6i-m). To reduce 3D migration speed without modifying the chemokine gradient, we treated ECs with low doses of Marimastat (1 mM) to reduce MMP activity. In similar fashion to decreased [S1P], Marimastat treatment decreased invasion speed and increased EC invasion as multicellular sprouts (Fig. 6i-m). Taken together, an interplay of soluble and physical microenvironmental cues regulate EC migration and proliferation, and a critical balance between these two basic cell functions is required for multicellular sprout invasion (Fig. 6n).

## DISCUSSION

EC migration and proliferation have been previously identified as key requirements for the formation of microvasculature in several models of angiogenesis^29,30^, but the relationship between these two fundamental cell functions has not been established. Furthermore, assessing the integrity of multicellular sprouts and resulting function of formed microvasculature has been challenging in previous *in vitro* model systems. In this work, we streamlined the fabrication of a multiplexed microfluidic device that recapitulates key aspects of 3D angiogenic sprouting and enables functional assessments of microvasculature fluidic connectivity and diffusive permeability. Tuning soluble and physical microenvironmental factors that affect EC migration and proliferation, we found that these two fundamental cell functions must be in balance to drive multicellular strand-like invasion of connected lumenized sprouts. Furthermore, microenvironmental conditions that balanced EC migration and proliferation yielded fluidically patent microvasculature with low diffusive permeability, two key traits of functional microvasculature. In stark contrast, imbalanced soluble or physical microenvironmental cues that elicited disproportionate migration and proliferation caused tip cells to break away from ensuing stalk cells; this resulted in disconnected ECs, blunt ended sprouts, fluidic leakiness, and high diffusive permeability.

While here we examined ECM density by tuning collagen concentration, our previous work demonstrates that ECM degradability also regulates the relationship between EC invasion speed and single vs. collective migration phenotypes, where highly degradable synthetic hydrogels increased EC invasion speed and invasion as single cells^27^. Of note, these synthetic matrices elicited limited EC proliferation; coupled with the findings presented here, insufficient proliferation may in part explain why multicellular sprouts failed to lumenize in those studies. Other cell functions that influence 3D cell migration efficiency such as proteolytic activity, cell-cell and cell-ECM adhesions, nuclear rigidity, and cytoskeletal contractility may also feed into whether cells migrate as single cells vs. collective multicellular strands^27,35–37^ (Fig. 6n). Cell migration mode is critical in both developmental and disease processes^38^. Cancer cells, for example, display a diversity of migratory phenotypes during metastasis including single cells or collective strands^37,39^. How microenvironmental cues regulate cell invasion mode and further, how invasion mode subsequently impacts metastatic efficiency are both important unanswered questions.

A balance of pro- and anti-angiogenic factors normally guide physiologic angiogenesis, but in many diseases an imbalance of soluble factors can dysregulate angiogenesis^8^. Prior to metastasis, primary tumors stimulate rapid angiogenesis to sustain the increasing metabolic demand of the growing tumor mass^40^. The tumor vasculature, however, is disorganized and hyperpermeable with heterogeneous EC populations and acellular gaps along the vessel wall^9,41,42^. In our model system, high [S1P] drove excessively high EC invasion speeds resulting in disconnected and highly permeable microvasculature, mimicking the rapid assembly of poor quality tumor vasculature. Indeed, elevated S1P levels have previously been observed in breast cancer murine models, where paracrine signaling from breast cancer cells secreting S1P enhances tumor angiogenesis and tumor burden^43,44^. Hyperpermeable, tortuous vasculature impairs proper blood flow, which hampers traditional therapeutic delivery and oxygen transport. Hypoxic conditions further reduce the susceptibility of the tumor to radio- and chemotherapy^45^. Given recent efforts to normalize vascular phenotype to better treat solid tumors, on-a-chip models integrating cancer cells with disorganized vasculature could provide a novel testbed for anti-cancer strategies focused on rescuing tumor microvasculature phenotype and function^46^. Based on the findings of our work, localized delivery of EC-targeting therapeutics that dually modulate invasion and proliferation may prevent or slow the progression of angiogenesis-mediated diseases.

Besides excessive angiogenesis, insufficient angiogenesis also contributes to disease progression. Seemingly paradoxical, both excessive and insufficient angiogenesis occur simultaneously in distinct organs of patients suffering from diabetes mellitus^10^. In the retina, the onset of EC proliferation delineates the transition between low (non-proliferative) and high (proliferative) grade diabetic retinopathy where rapid angiogenesis forms hyperpermeable microvasculature that contributes to eventual blindness^47^. Many of these same patients also present with impaired angiogenesis in the lower extremities, which inhibits wound healing and contributes to chronic diabetic foot ulcers^10^. A close examination of EC proliferation and invasion during angiogenesis through the lens of altered soluble and physical microenvironmental factors in diabetic tissues could help inform new strategies to halt or promote angiogenesis in the retina or skin, respectively. Outside of diabetes, there are currently no available therapies to address insufficient angiogenesis in ischemic conditions such as critical limb and cardiac ischemia^12^. Towards the design of vascularized tissue transplants to treat such conditions, this work emphasizes a need for balanced EC migration and proliferation for functional angiogenesis.

The vascularization of large bioengineered tissue constructs at organ relevant scales remains an outstanding challenge in the tissue engineering and regenerative medicine community^11^. Recent advances in 3D printing technologies enable exquisite control over arteriole and venule scale microchannels in synthetic hydrogel matrices, with such constructs subsequently endothelialized by flow-through seeding^48^. However, current 3D bioprinting approaches cannot achieve the 5-25 µm diameter length scale relevant to capillaries^2,33,49^. Furthermore, given that capillaries are narrower than fluid-suspended ECs (20-30 µm), flow-through seeding would prove difficult. Therefore, an integrated approach of 3D printed arteriole/venule-scale vasculature followed by controlled angiogenesis to elaborate the smallest scale capillaries may hold the most promise for generating functional hierarchical microvascular beds with potential for surgical integration^50^. However, a key challenge will be identifying hydrogel properties that enable high fidelity 3D printing while supporting angiogenesis, as several reports have indicated EC mechanosensitivity during migration, proliferation, and sprouting^15,21,27,51^. Overall, continued efforts to carefully dissect how architectural and mechanical attributes of the surrounding 3D space influence angiogenesis are critical (Fig. 6n). Here, we utilized collagen hydrogels to model the collagenous stroma where angiogenesis typically occurs. However, it is challenging to orthogonally tune material properties such as stiffness, degradability, and ligand density in natural materials such as collagen^52^. Thus, the continued development of synthetic hydrogels and their integration with microfluidic devices will be critical to shedding deeper insight into how specific aspects of the ECM regulate angiogenesis. As many commonly utilized synthetic hydrogels lack the fibrous architecture of native tissues, recent developments in fiber-reinforced synthetic hydrogels may provide novel insights into how fibrous cues regulate EC sprouting morphology^53^. The information gleaned from such studies would provide rich data sets to inform computational models of angiogenic sprouting and help in identifying mechanochemical design parameters that optimize implant vascularization for regenerative tissue therapies^54^. A multi-disciplinary approach combining on-a-chip platforms, synthetic biomaterials, imaging of live cell molecular reporters, and computational modeling would enable prediction and control of angiogenesis across a diversity of tissue environments, essential to elucidating mechanisms of disease progression, designing therapeutics to normalize vasculature, and engineering vascularized biomaterial implants.

## METHODS

### Reagents

All reagents were purchased from Sigma-Aldrich and used as received, unless otherwise stated.

### Microfluidic device fabrication

3D printed moulds were designed in AutoCAD and printed via stereolithography from Protolabs (Maple Plain, MN). Polydimethylsiloxane (PDMS, 1:10 crosslinker:base ratio) devices were replica casted from 3D printed moulds, cleaned with isopropyl alcohol and ethanol, and bonded to glass coverslips with a plasma etcher. Devices were treated with 0.01% (w/v) poly-l-lysine and 0.5% (w/v) L-glutaraldehyde sequentially for 1 hour each to promote ECM attachment to the PDMS housing, thus preventing hydrogel compaction from cell-generated forces. 300 µm stainless steel acupuncture needles (Lhasa OMS, Weymouth, MA) were inserted into each device and sterilized. Type I rat tail collagen (Corning, Corning, NY) was prepared as in Doyle^55^, injected into each device, and polymerized around each set of needles for 30 minutes at 37°C. Collagen hydrogels were hydrated in EGM2 for 2 hours and needles were removed to form 3D hollow channels fully embedded within collagen, positioned 400 µm away from PDMS and glass boundaries.

### Device cell seeding and culture

Human umbilical vein endothelial cells (HUVEC, Lonza, Switzerland) were cultured in endothelial growth media (EGM2, Lonza). HUVECs were passaged upon achieving confluency at a 1:4 ratio and used in studies from passages 4 to 9. A 20 µl solution of suspended HUVECs was added to one reservoir of the endothelial channel and inverted to allow cell attachment to the top half of the channel, followed by a second seeding with the device upright to allow cell attachment to the bottom half of the channel. HUVEC solution density was varied with collagen density as attachment efficiency was dependent on collagen density (1.5 M/ml for 2 mg ml^−1^, 2 M/ml for 3 mg ml^−1^ and 5 M/ml for 6 mg ml^−1^). HUVEC seeding densities were determined experimentally to achieve parent vessels with consistent cell densities across each collagen density (Supplemental Fig. 1c). HUVECs reached confluency and self-assembled into stable parent vessels over 24 hours. Media and chemokines were refreshed every 24 hours and devices were cultured with continual reciprocating flow. To inhibit cell proliferation, cells were treated with 40 µg ml^−1^ mitomycin C for 2 hours. To inhibit S1P receptor, cells were treated with 100 nM FTY720 for 24 hours.

### Lentivirus production

cDNA for pLenti CMV GFP Puro (658-5) was a gift from Eric Campeau and Paul Kaufman (Addgene plasmid #17448^56^). To generate lentivirus, plasmids were co-transfected with pCMV-VSVG (a gift from Bob Weinberg, Addgene plasmid #8454), pMDLg/pRRE, and pRSV-REC (gifts from Didier Trono, Addgene plasmid #12251 and #12253^57,58^) in 293T cells using the calcium phosphate precipitation method. Viral supernatants were collected after 48h, concentrated with PEG-it™ (System Biosciences, Palo Alto, CA) following the manufacturer’s protocol, filtered through a 0.45 µm filter (ThermoFisher Scientific Nalgene, Waltham, MA), and stored at −80°C. Viral titer was determined by serial dilution and infection of HUVECs in the presence of 10 µg ml^−1^ polybrene (Santa Cruz Biotechnology, Dallas, TX). Titers yielding maximal expression without cell death or detectable impact on cell proliferation or morphology were selected for studies.

### Fluorescent staining

Samples were fixed with 4% paraformaldehyde and permeabilized with a PBS solution containing Triton X-100 (5% v/v), sucrose (10% w/v), and magnesium chloride (0.6% w/v) for 1 hour each at room temperature. AlexaFluor 488 phalloidin (Life Technologies, Carlsbad, CA) was utilized to visualize F-actin. 4’, 6-diamidino-2-phenylindole (Sigma Aldrich) was utilized to visualize cell nucleus. For proliferation studies, EdU labelling was performed following the manufacturer’s protocol (ClickIT EdU, Life Technologies). DyLight 649 labelled Ulex Europaeus Agglutinin-1 (UEA, Vector Labs, Burlingame, CA) was utilized to visualize endothelial cell morphology in samples stained with EdU due to EdU ClickIT incompatibility with phalloidin staining. To visualize VE-cadherin, samples were sequentially blocked in bovine serum albumin (0.3% w/v), incubated with primary mouse monoclonal anti-VE-cadherin (1:1000, Santa Cruz Biotechnology), and incubated with secondary AlexaFluor 647 goat anti-mouse IgG (H+L) (1:1000, Life Technologies) each for 1 hour at room temperature. Fluorescently labelled collagen hydrogels were prepared as in Doyle^55^.

### Microscopy and image analysis

Fluorescent images were captured on a Zeiss LSM800 confocal microscope. Parent vessel endothelial cell density and EdU proliferation was quantified by counting DAPI and EdU positive cell nuclei. Invasion depth was quantified as the distance from the parent vessel edge to the tip cell and measured in FIJI. Invasion depth measurements were performed at 100 µm intervals along the parent vessel (Supplemental Fig. 2). Leading edge single cells were quantified as the number of single cells in the leading 150 µm front of cell invasion (Supplemental Fig. 2). Sprouts were quantified as the number of connected multicellular sprouts (parent vessel edge to tip cell) with a length greater than half the maximum invasion depth per condition (Supplemental Fig. 2). Sprout diameter measurements were performed in FIJI; diameter measurements smaller than the width of a cell nuclei (10 µm) were not included in the analysis. To assess fluidic connectivity and diffusive permeability, endothelial cells were first allowed to invade and reach the chemokine channel over 10-14 day culture. Diffusive permeability was quantified as in Polacheck et al.^59^ Fluorescent dextran (70 kDa Texas Red, Thermo Fisher) was incorporated into EGM2 media at 12.5 ug ml^−1^ and dextran diffusion was imaged at 1 second intervals to measure the flux of dextran from neovessels into the ECM. The resulting diffusion profile was fitted to a dynamic mass-conservation equation as in Adamson et al.^60^ with the diffusive-permeability coefficient (*P*_D_) defined by *J* = *P*_D_(*c*_vessel_−*c*_ECM_), where *J* is the mass flux of dextran, *c*_vessel_ is the concentration of dextran in the vessel, and *c*_ECM_ is the concentration of dextran in the perivascular ECM.

### Statistics

Statistical significance was determined by one-way analysis of variance (ANOVA) or two-sided student’s t-test where appropriate, with significance indicated by p<0.05. Sample size is indicated within corresponding figure legends and all data are presented as mean ± standard deviation.

## Data availability

The data that support the findings of this study are available from the corresponding author upon reasonable request.

## Acknowledgements

This work was supported in part by the National Institutes of Health (HL124322). W.Y.W acknowledges financial support from the University of Michigan Rackham Merit Fellowship and the National Science Foundation Graduate Research Fellowship Program (DGE1256260).

## Author contributions

W.Y.W. and B.M.B. designed the experiments. W.Y.W, D.L., and E.H.J. conducted experiments and analyzed the data. W.J.P. performed permeability analysis. W.Y.W. and B.M.B. wrote the manuscript. All authors reviewed the manuscript.

## References

1. Huxley, V. & Rumbaut, R. The Microvasculature As A Dynamic Regulator Of Volume And Solute Exchange. Clin. Exp. Pharmacol. Physiol. 27, 847–854 (2000).

2. Aird, W. C. Spatial and temporal dynamics of the endothelium. Journal of Thrombosis and Haemostasis 3, 1392–1406 (2005).

3. Johnson, J. M., Minson, C. T. & Kellogg, D. L. Cutaneous vasodilator and vasoconstrictor mechanisms in temperature regulation. Compr. Physiol. 4, 33–89 (2014).

4. Tonnesen, M. G., Feng, X. & Clark, R. A. F. F. Angiogenesis in wound healing. J. Investig. Dermatology Symp. Proc. 5, 40–46 (2000).

5. Fraisl, P., Mazzone, M., Schmidt, T. & Carmeliet, P. Regulation of Angiogenesis by Oxygen and Metabolism. Dev. Cell 16, 167–179 (2009).

6. Wimmer, R. A. et al. Human blood vessel organoids as a model of diabetic vasculopathy. Nature 565, 505–510 (2019).

7. Carmeliet, P. Angiogenesis in life, disease and medicine. Nature 438, 932–936 (2005).

8. Carmeliet, P. & Jain, R. K. Angiogenesis in cancer and other diseases. Nature 407, 249–257 (2000).

9. Siemann, D. W. The unique characteristics of tumor vasculature and preclinical evidence for its selective disruption by Tumor-Vascular Disrupting Agents. Cancer Treat. Rev. 37, 63–74 (2011).

10. Martin, A., Komada, M. R. & Sane, D. C. Abnormal angiogenesis in diabetes mellitus. Med. Res. Rev. 23, 117–145 (2003).

11. Novosel, E. C., Kleinhans, C. & Kluger, P. J. Vascularization is the key challenge in tissue engineering. Adv. Drug Deliv. Rev. 63, 300–311 (2011).

12. Losordo, D. W. & Dimmeler, S. Therapeutic angiogenesis and vasculogenesis for ischemic disease. Part II: Cell-based therapies. Circulation 109, 2692–2697 (2004).

13. Francavilla, C., Maddaluno, L. & Cavallaro, U. The functional role of cell adhesion molecules in tumor angiogenesis. Semin. Cancer Biol. 19, 298–309 (2009).

14. Potente, M., Gerhardt, H. & Carmeliet, P. Basic and Therapeutic Aspects of Angiogenesis. Cell 146, 873–887 (2011).

15. Crosby, C. O. & Zoldan, J. Mimicking the physical cues of the ECM in angiogenic biomaterials. Regen. Biomater. 6, 61–73 (2019).

16. Nowak-Sliwinska, P. et al. Consensus guidelines for the use and interpretation of angiogenesis assays. Angiogenesis 21, 425–532 (2018).

17. Hayer, A. et al. Engulfed cadherin fingers are polarized junctional structures between collectively migrating endothelial cells. Nat. Cell Biol. 18, 1311–1323 (2016).

18. Baker, B. M. & Chen, C. S. Deconstructing the third dimension – how 3D culture microenvironments alter cellular cues. J. Cell Sci. 125, 3015–3024 (2012).

19. Gerhardt, H. et al. VEGF guides angiogenic sprouting utilizing endothelial tip cell filopodia. J. Cell Biol. 161, 1163–1177 (2003).

20. Jakobsson, L. et al. Endothelial cells dynamically compete for the tip cell position during angiogenic sprouting. Nat. Cell Biol. 12, 943–953 (2010).

21. Bordeleau, F. et al. Matrix stiffening promotes a tumor vasculature phenotype. Proc. Natl. Acad. Sci. 114, 492–497 (2017).

22. Chen, M. B. et al. On-chip human microvasculature assay for visualization and quantification of tumor cell extravasation dynamics. Nat. Protoc. 12, 865–880 (2017).

23. Polacheck, W. J. et al. A non-canonical Notch complex regulates adherens junctions and vascular barrier function. Nature 552, 258–262 (2017).

24. Akbari, E., Spychalski, G. B. & Song, J. W. Microfluidic approaches to the study of angiogenesis and the microcirculation. Microcirculation 24, e12363 (2017).

25. Alimperti, S. et al. Three-dimensional biomimetic vascular model reveals a RhoA, Rac1, and N-cadherin balance in mural cell–endothelial cell-regulated barrier function. Proc. Natl. Acad. Sci. 114, 8758–8763 (2017).

26. Nguyen, D.-H. T. et al. Biomimetic model to reconstitute angiogenic sprouting morphogenesis in vitro. Proc. Natl. Acad. Sci. U. S. A. 110, 6712–6717 (2013).

27. Trappmann, B. et al. Matrix degradability controls multicellularity of 3D cell migration. Nat. Commun. 8, 371 (2017).

28. Paik, J. H., Chae Ss, S., Lee, M. J., Thangada, S. & Hla, T. Sphingosine 1-phosphate-induced endothelial cell migration requires the expression of EDG-1 and EDG-3 receptors and Rho-dependent activation of alpha vbeta3- and beta1-containing integrins. J. Biol. Chem. 276, 11830–7 (2001).

29. Ausprunk, D. H. & Folkman, J. Migration and proliferation of endothelial cells in preformed and newly formed blood vessels during tumor angiogenesis. Microvasc. Res. 14, 53–65 (1977).

30. Pontes-Quero, S. et al. High mitogenic stimulation arrests angiogenesis. Nat. Commun. 10, 2016 (2019).

31. Cross, V. L. et al. Dense type I collagen matrices that support cellular remodeling and microfabrication for studies of tumor angiogenesis and vasculogenesis in vitro. Biomaterials 31, 8596–8607 (2010).

32. Osaki, T. et al. Acceleration of Vascular Sprouting from Fabricated Perfusable Vascular-Like Structures. PLoS One 10, e0123735 (2015).

33. Nunes, S. S. et al. Implanted microvessels progress through distinct neovascularization phenotypes. Microvasc. Res. 79, 10–20 (2010).

34. Adamson, R. H., Huxley, V. H. & Curry, F. E. Single capillary permeability to proteins having similar size but different charge. Am. J. Physiol. Circ. Physiol. 254, H304–H312 (1988).

35. Krause, M. et al. Cell migration through 3D confining pores: speed accelerations by deformation and recoil of the nucleus. Transactions B: Biological Sciences 374, (2019).

36. Fraley, S. I. et al. A distinctive role for focal adhesion proteins in three-dimensional cell motility. Nat. Cell Biol. 12, 598–604 (2010).

37. Wang, W. Y., Davidson, C. D., Lin, D. & Baker, B. M. Actomyosin contractility-dependent matrix stretch and recoil induces rapid cell migration. Nat. Commun. 10, 1186 (2019).

38. Friedl, P. & Gilmour, D. Collective cell migration in morphogenesis, regeneration and cancer. Nat. Publ. Gr. 10, (2009).

39. Wolf, K. et al. Multi-step pericellular proteolysis controls the transition from individual to collective cancer cell invasion. Nat. Cell Biol. 9, 893–904 (2007).

40. Almog, N. et al. Transcriptional switch of dormant tumors to fast-growing angiogenic phenotype. Cancer Res. 69, 836–844 (2009).

41. Hashizume, H. et al. Openings between defective endothelial cells explain tumor vessel leakiness. Am. J. Pathol. 156, 1363–1380 (2000).

42. Di Tomaso, E. et al. Mosaic tumor vessels: Cellular basis and ultrastructure of focal regions lacking endothelial cell markers. Cancer Res. 65, 5740–5749 (2005).

43. Nagahashi, M. et al. Sphingosine-1-phosphate produced by sphingosine kinase 1 promotes breast cancer progression by stimulating angiogenesis and lymphangiogenesis. Cancer Res. 72, 726–735 (2012).

44. Takabe, K. et al. Estradiol induces export of sphingosine 1-phosphate from breast cancer cells via ABCC1 and ABCG2. 285, 10477–10486 (2010).

45. Declerck, K. & Elble, R. C. The role of hypoxia and acidosis in promoting metastasis and resistance to chemotherapy. Frontiers in Bioscience 15, 213–225 (2010).

46. Jain, R. K. Normalization of tumor vasculature: an emerging concept in antiangiogenic therapy. Science 307, 58–62 (2005).

47. Durham, J. T. & Herman, I. M. Microvascular modifications in diabetic retinopathy. Current Diabetes Reports 11, 253–264 (2011).

48. Grigoryan, B. et al. Multivascular networks and functional intravascular topologies within biocompatible hydrogels. Science 364, 458–464 (2019).

49. Gong, H., Beauchamp, M., Perry, S., Woolley, A. T. & Nordin, G. P. Optical approach to resin formulation for 3D printed microfluidics. RSC Adv. 5, 106621–106632 (2015).

50. Mirabella, T. et al. 3D-printed vascular networks direct therapeutic angiogenesis in ischaemia. Nat. Biomed. Eng. 1, 0083 (2017).

51. Davidson, C. D., Wang, W. Y., Zaimi, I., Jayco, D. K. P. & Baker, B. M. Cell force-mediated matrix reorganization underlies multicellular network assembly. Sci. Rep. 9, 12 (2019).

52. Li, L., Eyckmans, J. & Chen, C. S. Designer biomaterials for mechanobiology. Nat. Mater. 16, 1164–1168 (2017).

53. Matera, D. L., Wang, W. Y., Smith, M. R., Shikanov, A. & Baker, B. M. Fiber Density Modulates Cell Spreading in 3D Interstitial Matrix Mimetics. ACS Biomater. Sci. Eng. 5, 2965–2975 (2019).

54. Heck, T. A. M., Vaeyens, M. M. & Van Oosterwyck, H. Computational Models of Sprouting Angiogenesis and Cell Migration: Towards Multiscale Mechanochemical Models of Angiogenesis. Math. Model. Nat. Phenom. 10, 108–141 (2015).

55. Doyle, A. D. Generation of 3D collagen gels with controlled diverse architectures. Curr. Protoc. Cell Biol. 2016, 10.20.1–10.20.16 (2016).

56. Campeau, E. et al. A versatile viral system for expression and depletion of proteins in mammalian cells. PLoS One 4, e6529(2009).

57. Stewart, S. A. et al. Lentivirus-delivered stable gene silencing by RNAi in primary cells Lentivirus-delivered stable gene silencing by RNAi in primary cells. Rna 9, 493–501 (2003).

58. Dull, T. et al. A third-generation lentivirus vector with a conditional packaging system. J. Virol. 72, 8463–71 (1998).

59. Polacheck, W. J., Kutys, M. L., Tefft, J. B. & Chen, C. S. Microfabricated blood vessels for modeling the vascular transport barrier. Nat. Protoc. 14, 1425–1454 (2019).

60. Adamson, R. H., Lenz, J. F. & Curry, F. E. Quantitative Laser Scanning Confocal Microscopy on Single Capillaries: Permeability Measurement. Microcirculation 1, 251–265 (1994).

