## Supplemental Material for "Functional angiogenesis requires microenvironmental cues balancing endothelial cell migration and proliferation"

### SUPPLEMENTARY MATERIAL

The Supplementary Material includes 4 Supplementary Figures.

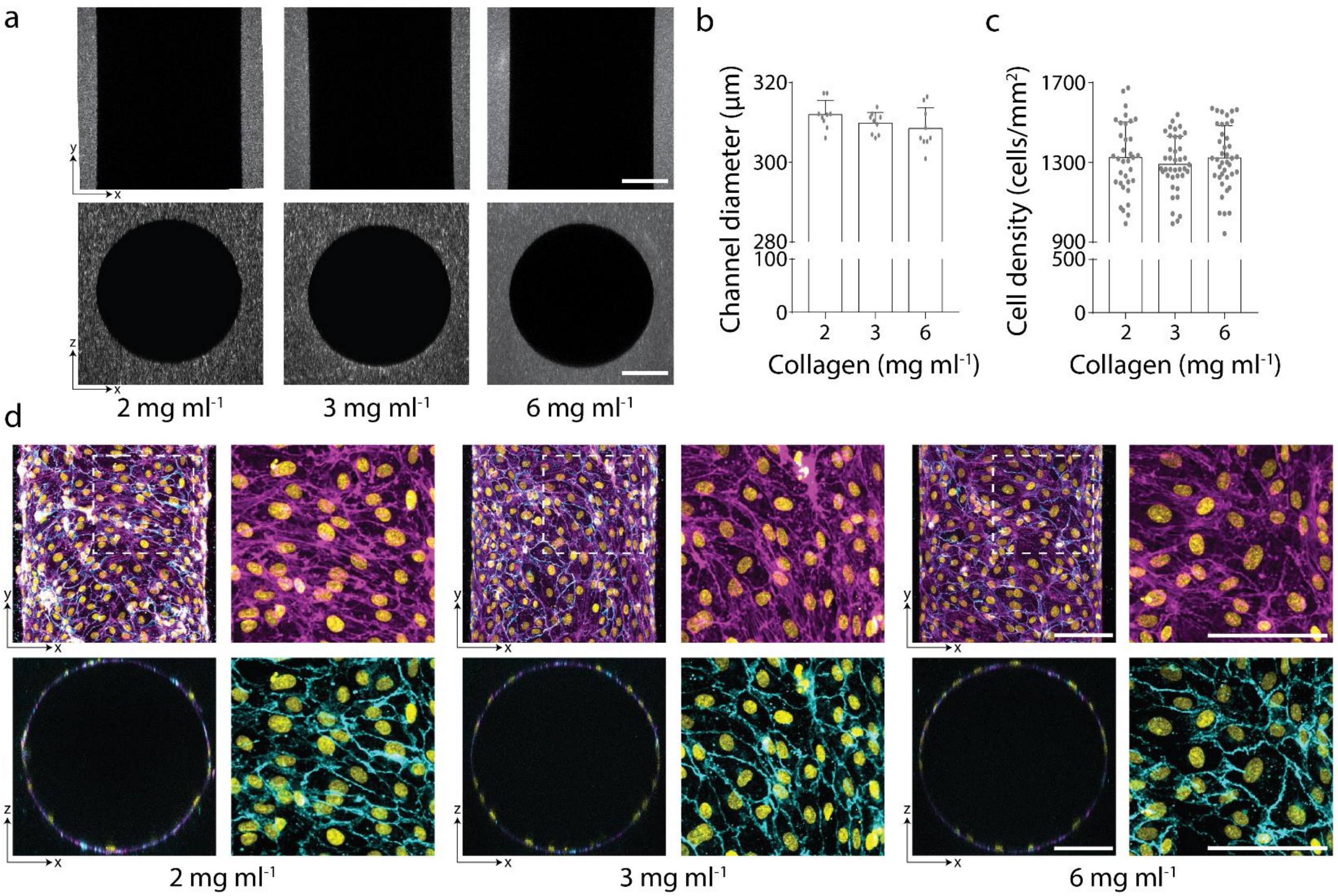


**Supplemental Figure 1 | Parent vessels with consistent diameter and cell density. a,** Representative images of x-y and x-z orthogonal views of 3D channels generated in indicated collagen density. **b-c,** Quantifications of channel diameter and parent vessel endothelial cell density with varying collagen concentration. All data presented as mean ± s.d.; * indicates a statically significant comparison with $P<0.05$ (one-way analysis of variance). $n \geq 8$ channels (**b**) and $n \geq 34$ vessel segments (each 400 μm length) (**c**) per condition. **d,** Representative images of x-y (max intensity projection) and x-z (single slice) orthogonal views of parent vessels generated with varying collagen concentration after 24-hour culture. Insets indicated with dashed white lines. Nucleus (yellow), F-actin (magenta), and VE-cadherin (cyan).

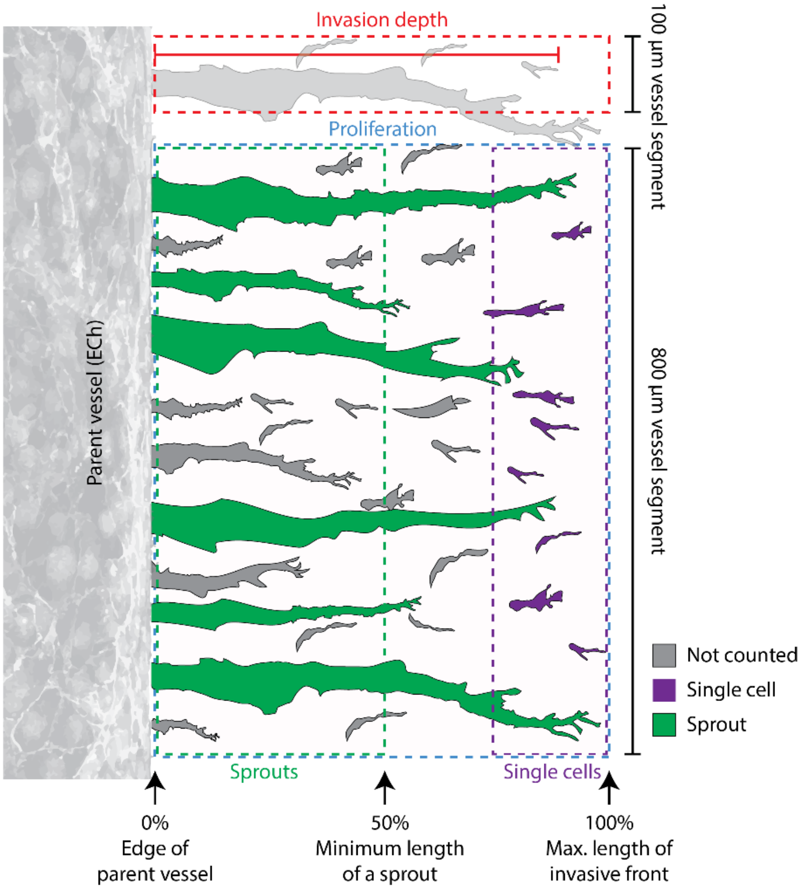


**Supplemental Figure 2 | Schematic of quantification metrics.** To quantify invasion depth (red), images were segmented into 100 µm segments along the parent vessel and measurements were taken from the ECh edge to the tip of the endothelial cell that invaded furthest within that segment. To quantify proliferation (blue), images were segmented into 800 µm segments along the parent vessel, and the percentage of $EdU^+$ nuclei was measured only in endothelial cells that had invaded into the extracellular matrix from the ECh edge to leading invasive front. To quantify leading edge single cells (purple), images were segmented into 800 µm segments along the parent vessel and isolated single cells were quantified within 150 µm of the leading invasive front. To quantify multicellular sprouts (green), images were segmented into 800 µm segments along the parent vessel, and quantified connected endothelial sprout structures (from ECh edge to sprout tip) with a minimum length of half the max invasion depth. Single cells and sprouts outside of these criteria were not included in the analysis (grey).

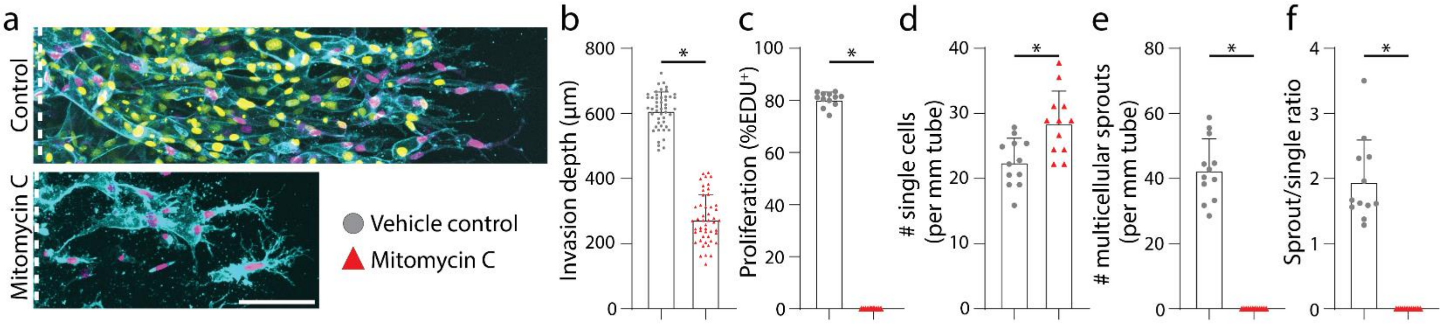


**Supplemental Figure 3 | Proliferation inhibition abrogates multicellular sprouting. a,** Representative images (max intensity projection) of endothelial cell invasion in response to mitomycin C proliferation inhibition with S250:P50 and 3mg $ml^{-1}$ collagen. **b-f,** Quantifications of invasion depth, proliferation, and morphology of invading endothelial cells as single cells or multicellular sprouts. All data presented as mean ± s.d.; * indicates a statically significant comparison with $P<0.05$ (two-tailed Student's t-test). For invasion depth analysis (**b**), n=48 vessel segments (each 100 µm length) per condition. For proliferation and migration mode analysis (**c-f**) n=12 vessel segments (each 800 µm length) per condition.

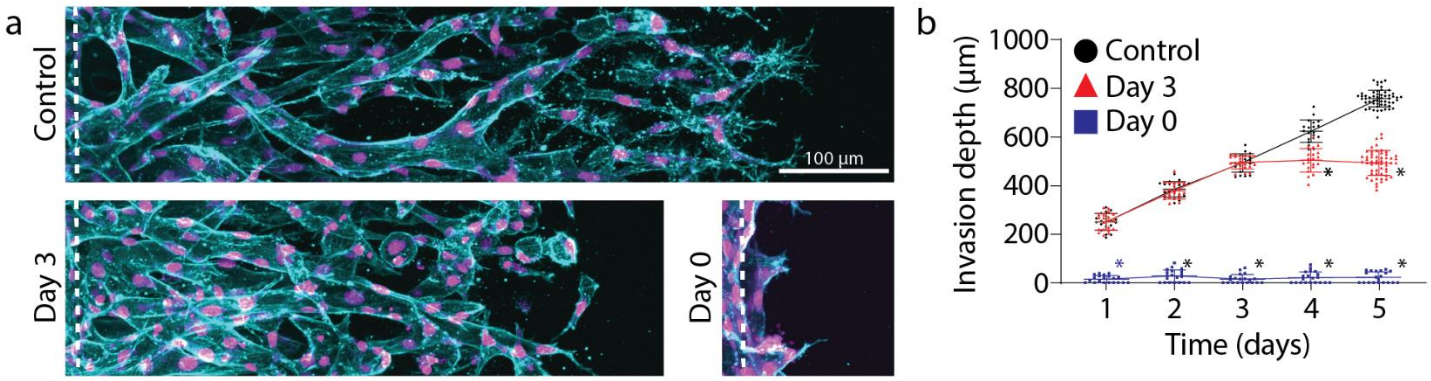


**Supplemental Figure 4 | S1P receptor inhibition abrogates S1P-driven EC invasion. a,** Representative images (max intensity projection) of invading endothelial cells in response to 100 nM FTY720 treatment with S250:P25 and 3mg ml$^{-1}$ collagen. Day 0 conditions were composed of ECs pre-treated with FTY720. F-actin (cyan), nucleus (magenta). **b,** Endothelial cell invasion depth over time in response to 100 nM FTY720 treatment. All data presented as mean ± s.d.; * indicates a statically significant comparison with $P<0.05$ (one-way analysis of variance). $n\geq20$ vessel segments (each 100 µm length) per condition.
